# How, when, and where relic DNA biases estimates of microbial diversity

**DOI:** 10.1101/131284

**Authors:** JT Lennon, ME Muscarella, SA Muscarella, BK Lehmkuhl

## Abstract

Extracellular or “relic” DNA is one of the largest pools of nucleic acids in the mbiosphere ^1,2^. Relic DNA can influence a number of important ecological and evolutionary processes, but it may also bias estimates of microbial abundance and diversity, which has implications for understanding environmental, engineered, and host-associated ecosystems. We developed models capturing the fundamental processes that regulate the size and composition of the relic DNA pools to identify scenarios leading to biased estimates of biodiversity. Our models predict that bias increases with relic DNA pool size, but only when the species abundance distributions (SAD) of relic and intact DNA are distinct from one another. We evaluated our model predictions by quantifying relic DNA and assessing its contribution to bacterial diversity using 16S rRNA gene sequences collected from different ecosystem types, including soil, sediment, water, and the mammalian gut. On average, relic DNA made up 33 % of the total bacterial DNA pool, but exceeded 80 % in some samples. Despite its abundance, relic DNA had no effect on estimates of taxonomic and phylogenetic diversity, even in ecosystems where processes such as the physical protection of relic DNA are common and predicted by our models to generate bias. Rather, our findings are consistent with the expectation that relic DNA sequences degrade in proportion to their abundance and therefore may contribute minimally to estimates of microbial diversity.

---

When microorganisms die, their DNA leaks into the surrounding environment. The fate of this relic DNA has important implications for evolutionary and ecological processes. For example, relic DNA can be taken up and incorporated into the genomes of some microorganisms via transformation, thereby serving as a reservoir of genetic information that can confer new traits and fitness benefits to distantly related taxa ^3^. In addition, relic DNA is a high-quality resource containing carbon, nitrogen, and phosphorus that is consumed by a diverse array of bacteria with consequences for microbial community structure and ecosystem processes ^1,4^.

Relic DNA also has the potential to bias cultivation-independent estimates of diversity, which are widely used for addressing questions concerning the assembly, biogeography, and functioning of microbial communities. Microbial DNA extracted from environmental and host-associated samples is not solely derived from metabolically active organisms ^5^. A large portion of the individuals in a microbial community is dormant or dead ^6,7^. Although nucleic acids can be temporarily retained in non-viable cells, DNA is ultimately released into the environment when individuals die from autolysis, senescence, viral infection, or predation ^8^. Together, these sources of mortality can create large pools of relic DNA ^2,9^. For example, there is an estimated 0.45 petagrams of relic DNA in global ocean sediments, which is 70-fold greater than the amount of DNA contained in intact cells from the same environments ^1^. If included in cultivation-independent approaches, relic DNA could distort our understanding of the ecological and evolutionary processes that regulate the distribution, abundance, and function of microbial taxa.

The processes that regulate relic DNA dynamics vary among ecosystems. In well-mixed microbial habitats, the residence time of relic DNA can be short owing to rapid rates of hydrolysis, oxidation, and UV-mediated damage ^2^. For example, in surface waters of freshwater and marine environments, the extracellular DNA pool turns over in less than one day ^10,11^ while plasmid DNA begins to degrade in just minutes ^12^. As a result, the size distribution of relic DNA is skewed towards small fragments ranging between 100 and 500 base pairs ^13,14^. In more structured microbial habitats, other factors contribute to the size and turnover of the relic DNA pool. For example, binding with inorganic and organic substances in soils and sediments reduces the rate of relic DNA degradation ^9^. Likewise, biofilms, aggregates, and outer membrane vesicles can protect relic DNA from hydrolytic enzymes ^8,15,16^. Alternatively, the heterogeneous distribution of microorganisms in structured habitats creates “hot spots” of metabolic activity^17^ that may increase relic DNA turnover. Collectively, these processes may help explain the accumulation of relic DNA in a range of ecosystems including ocean sediments ^1^, the built environment^18^, and the mammalian gastrointestinal tract ^19^.

To date, the documented effects of relic DNA on estimates of diversity are idiosyncratic. Even in samples with large amounts of relic DNA, bias can be non-existent or substantial, and can lead to the overestimation or underestimation of diversity ^20,21^. Such observations reflect the need for a more mechanistic understanding of relic DNA dynamics so that microbial communities from different ecosystems can be consistently and accurately characterized. We address this issue by developing a theoretical framework that considers the processes regulating the size and turnover of the relic DNA pool. We emphasize that the direction and magnitude of bias is influenced by sampling from a joint species abundance distribution (SAD) consisting of sequences from the relic and intact DNA pools. We evaluated our models by quantifying the contribution of relic DNA to the abundance and diversity of bacterial communities in ecosystem types that are known to have contrasting relic DNA dynamics.

## RESULTS

We developed an interrelated set of models to identify conditions where relic DNA leads to biased estimates of microbial diversity. We began with a conceptual model representing the dynamics of intact and relic DNA (Fig. 1a). The size and composition of the intact DNA pool reflects sequences contained in viable bacteria belonging to different species, which are influenced by births, deaths, and immigration. The size and composition of the relic DNA pool is determined by the input of sequences associated with the death of bacteria from the intact DNA pool and losses associated with the degradation of relic DNA sequences. With this framework established, we developed a sampling model to explore how diversity estimates are affected when draws come from a joint distribution of sequences derived from the intact and relic pools (see Methods). We found that a sufficiently large relic DNA pool is required, but not sufficient to create bias in estimates of species richness, the number of taxa in a sample (Fig. 1b). Critically, there must also be differences between the species abundance distributions (SAD) of the intact and relic DNA pools for bias to arise. When the relic DNA pool has a more even SAD, sampling from the total pool (intact + relic) leads to overestimation of true richness (Fig. 1b). Conversely, when the relic DNA pool has a less even SAD, sampling from the total pool leads to underestimation of true richness (Fig. 1b). Last, we developed a process-based model to identify conditions that lead to relic DNA bias. We determined that relic DNA pool size reaches a stable equilibrium when *R* = *m I* / *d*, where *R* is the size (number of sequences) of the relic DNA pool, *m* is the per capita mortality rate of bacteria in the intact DNA pool, *I* is the size (number of sequences) of the intact DNA pool, and *d* is the per capita degradation rate of the relic DNA sequences (see Methods). From this, it can be shown that *R* increases with the residence time (τ = *R* / *d)* of the relic DNA pool. We also used the process-based model to explore different scenarios that influence the SAD of the relic DNA pool. First, we assumed that the rate at which relic DNA sequences degrade is equivalent among species. Under this neutral expectation, the shapes of relic SAD and intact SAD are nearly identical. No bias arises under these conditions, regardless of the relic DNA pool size (Fig. 1c). Next, we simulated protection by reducing the degradation rates of relic DNA sequences belonging to randomly selected species (see Methods). This created a less even relic SAD, resulting in the underestimation of richness in the total DNA pool (Fig. 1c). Finally, we considered the scenario where there are “hot spots” of relic DNA degradation. When simulating accelerated degradation rates for abundant relic DNA sequences, there was a more even relic SAD that resulted in the overestimation of richness in the total DNA pool (Fig. 1c).

**Figure 1.**
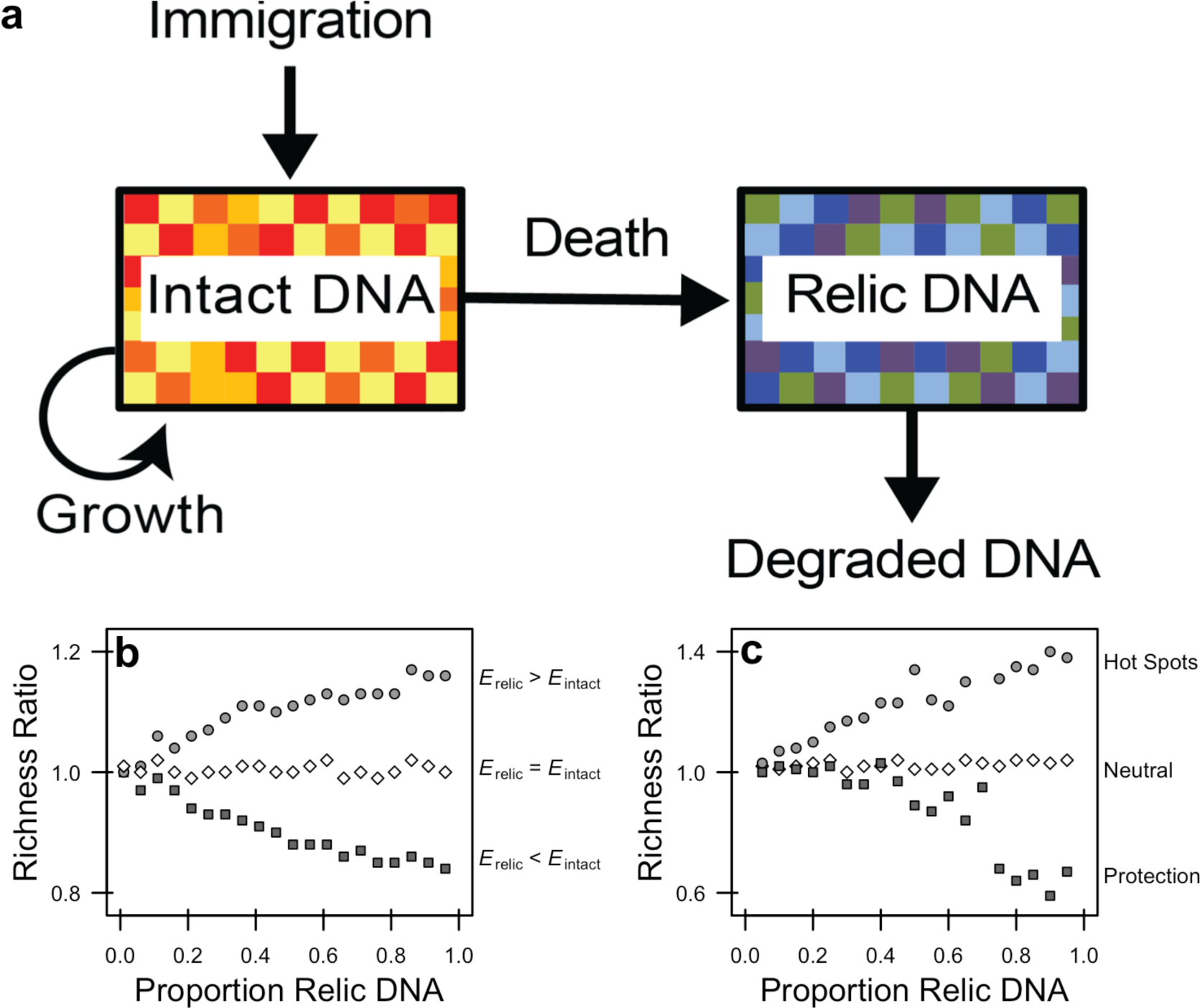
Modeling relic DNA dynamics. **a** The amount of relic DNA in a microbial environment is determined by inputs associated with the mortality of viable individuals with intact DNA and by losses associated with the degradation of relic DNA. If the diversity of sequences contained in the relic DNA pool is sufficiently different from that in the intact DNA pool, then relic DNA may bias estimates of microbial biodiversity (as indicated by different colored boxes) when sampling from the total (intact + relic) DNA pool. **b** We developed a sampling-based simulation model to explore the effects of mixing intact and relic DNA on estimates of diversity. We populated intact and relic communities with individuals from a lognormal species abundance distribution (SAD). We altered the diversity of the relic community by changing the scale parameter of the lognormal distribution describing the SAD. We then sampled and mixed the intact and relic communities so that the relic contribution to total community ranged from 0.01 to 0.96.**c** To gain mechanistic insight into how bias arises, we created a stochastic process-based model that captures features that influence relic DNA dynamics, including: immigration, birth, death, degradation (**a**). We simulated a range of degradation rates to achieve relic DNA pool sizes with proportions ranging between 0.05 and 0.95. To explore how degradation alters the SAD of the relic community, we explored three scenarios. First, we simulated a neutral scenario where relic DNA sequences produced by different species degrade at the same rate. Second, we simulated conditions where physical, chemical, or biological processes slow the degradation rate of relic DNA belonging to some species via protection. Third, we simulated “hot spots” where there more abundant relic DNA sequences experience faster rates of relic DNA degradation, a condition that may arise in structured habitats where there are patchy distributions of individuals and their metabolic products (i.e., enzymes). We ran simulations for 10,000 time steps and then sampled the intact and relic communities. To quantify bias in diversity (**b, c**), we calculated “richness ratios” which reflect the number of species in the total DNA pool (intact + relic) divided by the number of species in the intact DNA pool in a simulation. When values richness ratios = 1, relic DNA has no effect on estimates of diversity; when richness ratios > 1, relic DNA overestimates true diversity; and when richness ratios < 1, relic DNA underestimates true diversity.

### Variation in relic DNA pool size

We collected samples from different ecosystem types (soil, sediment, water, and mammalian guts) that are known to vary in relic DNA turnover rates ^13,22^. Using quantitative PCR with 16S rRNA primers, we quantified intact DNA after treating an aliquot of sample with DNase to remove relic DNA. In addition, we quantified total DNA (intact + relic) pool size from a control aliquot of sample that was not treated with DNase (see Methods and Fig. S2). Consistent with previous reports ^20,21,23^, relic DNA accounted for a substantial but variable portion of the total bacterial DNA pool. The proportion of relic DNA was normally distributed (mean ± sd: 0.33 ± 0.218) and ranged from 0 to 0.83 across 34 samples obtained from different ecosystem types (Fig. 2). Even though host and environmental features associated with different habitats are thought to influence relic DNA dynamics, there was only a marginally significant effect of ecosystem type on the proportion of relic DNA (one-way ANOVA, F_3_, _30_ = 2.43, *P* = 0.084) ^2,20^ which likely reflected the contrast between soil and water samples (Tukey’s HSD, adjusted *P* = 0.058). Nevertheless, the large average pool size and range in relic DNA across samples provided us with the opportunity to explore features of our model, specifically the magnitude and direction of bias that relic DNA should have on estimates of microbial diversity.

**Figure 2.**
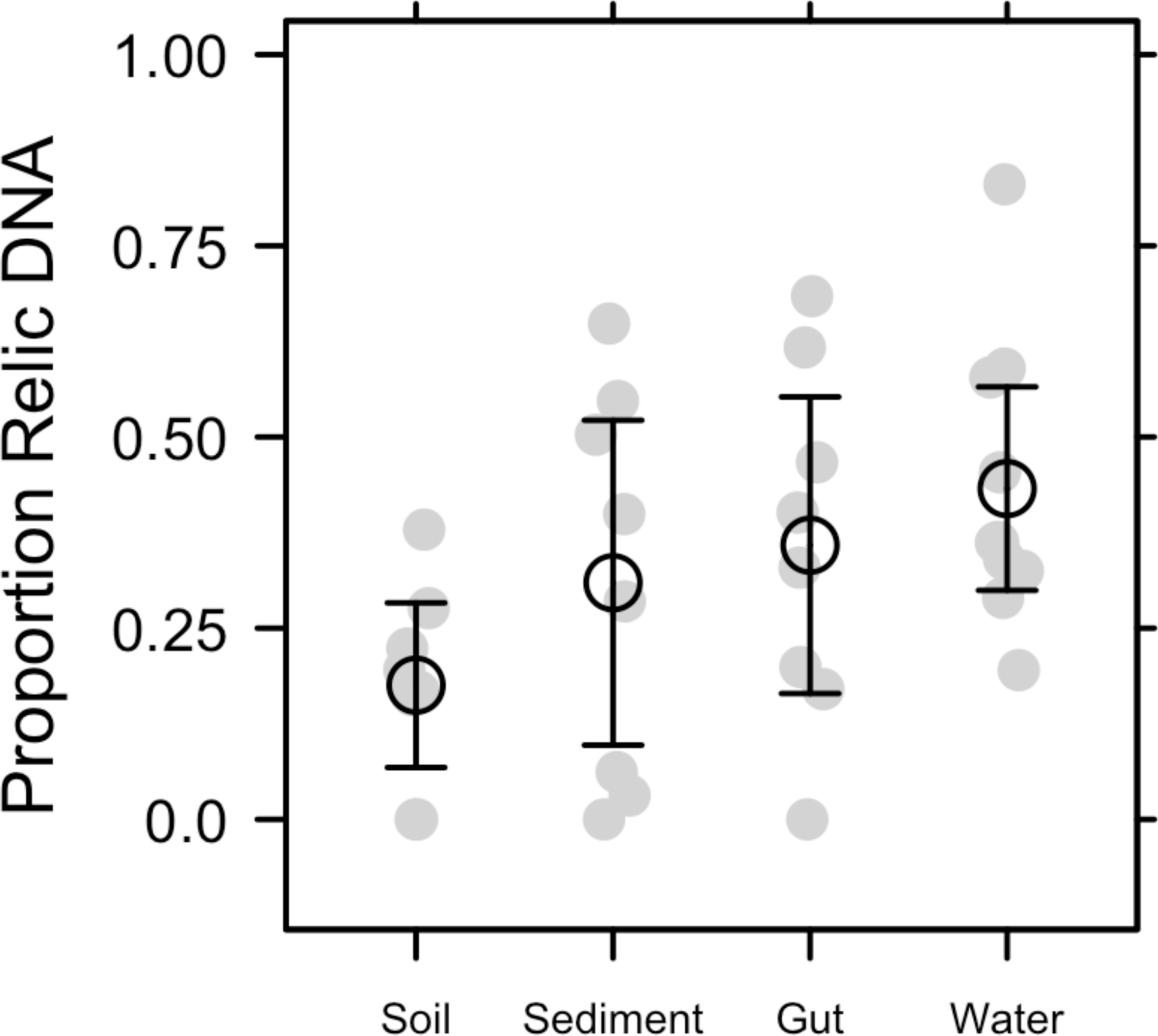
Proportion of bacterial relic DNA in different ecosystem types. We quantified the amount of intact DNA in a sample after removing relic DNA with a DNase treatment. We then estimated the proportion of relic DNA as 1 - (intact DNA / total DNA), where the total DNA concentration was quantified without DNase treatment. Relic DNA constituted an appreciable portion of the total DNA pool, but was not affected by the ecosystem type from which the sample was collected (gut, soil, sediment, and water). Grey symbols are the observed data and black symbols represent means ± 95% confidence intervals.

### Magnitude and direction of bias on microbial diversity

We sequenced 16S rRNA genes from the intact and total DNA pools to test whether or not relic DNA biases estimates of within-sample (alpha) bacterial diversity. Despite accounting for a substantial portion of the total DNA (Fig. 2), relic DNA had no effect on estimates of richness, evenness, or phylogenetic diversity (PD) (Table S1). We expressed each of these alpha-diversity metrics on a per-sample basis as a ratio (total / intact), where values > 1 represent overestimation bias and values < 1 represent underestimation bias. Relic DNA had no effect on the diversity ratios based on the observation that the 95 % confidence intervals overlapped with 1.0. Furthermore, the 95 % confidence intervals of the diversity ratios overlapped across ecosystem types, indicating that the contribution of relic DNA to all measures of diversity was generally low irrespective of the microbial habitat. Finally, using indicator variable multiple regression, we evaluated whether the magnitude of bias increased with relic DNA pool size, a predicted outcome from some of our simulations (Fig. 1b, c). For all diversity ratios (richness, evenness, and PD), the slopes were not different from zero (*P* ≥ 0.65), the intercepts were not different from 1.0 (*P* ≥ 0.65), and the estimates were not affected by ecosystem type (*P* ≥ 0.26) (Fig. S5). Moreover, only a very small amount of variation (R^2^ ≤ 0.01) was explained by relic DNA and ecosystem type in the regression models.

### Contribution of relic DNA to community composition

Based on 16S rRNA gene sequencing of the intact and total DNA pools, relic DNA had no effect on the compositional (beta) diversity of bacterial communities within or across ecosystem types. First, the intact and total DNA pools were significantly and very highly correlated with one another when we performed a taxonomically based (Bray-Curtis distances) Mantel test (*P* = 0.001, *r* = 0.959) and a phylogenetically based (UniFrac distances) Mantel text (*P* = 0.001, *r* = 0.996). Second, we tested for the effect of relic DNA on bacterial community composition for the intact and total DNA pools using a modified beta-dispersion metric^24^. Specifically, we calculated centroid distance ratios, which directly compared the dispersion between pairs of DNase-treated and DNase-control subsamples (see Methods and Fig. S7). With this approach, if the centroid distance ratio was > 1, we concluded that relic DNA inflated beta-diversity; if the distance ratio was < 1, we concluded that relic DNA homogenized beta-diversity. We found that relic DNA had no effect on the centroid distance ratios based on the observation that the 95 % confidence intervals overlapped with 1.0 (Figs. 4, S8). Furthermore, the 95 % confidence intervals for the centroid distance ratios overlapped across ecosystem types, indicating that the contribution of relic DNA to taxonomic and phylogenetic beta-diversity was low irrespective of the microbial habitat. Last, we used indicator variable multiple regression to test the prediction that bias in estimates of community composition would increase with increasing relic DNA pool size (see Fig. S1). The proportion of relic DNA in a sample had no effect on the slopes for centroid distance ratios regardless of whether they were calculated using taxonomic (*P* = 0.63) or phylogenetic (*P* = 0.59) distance matrices. Moreover, the intercepts for these regression relationships were not different from 1.0 (*P* ≥ 0.74) and the amount of explained variation was low (R^2^ ≤ 0.09) suggesting that the overall effect of relic DNA on beta diversity was negligible (Fig. S8). The parameters estimates from the regression analyses were not affected by ecosystems type (*P* ≥ 0.29).

**Figure 4.**
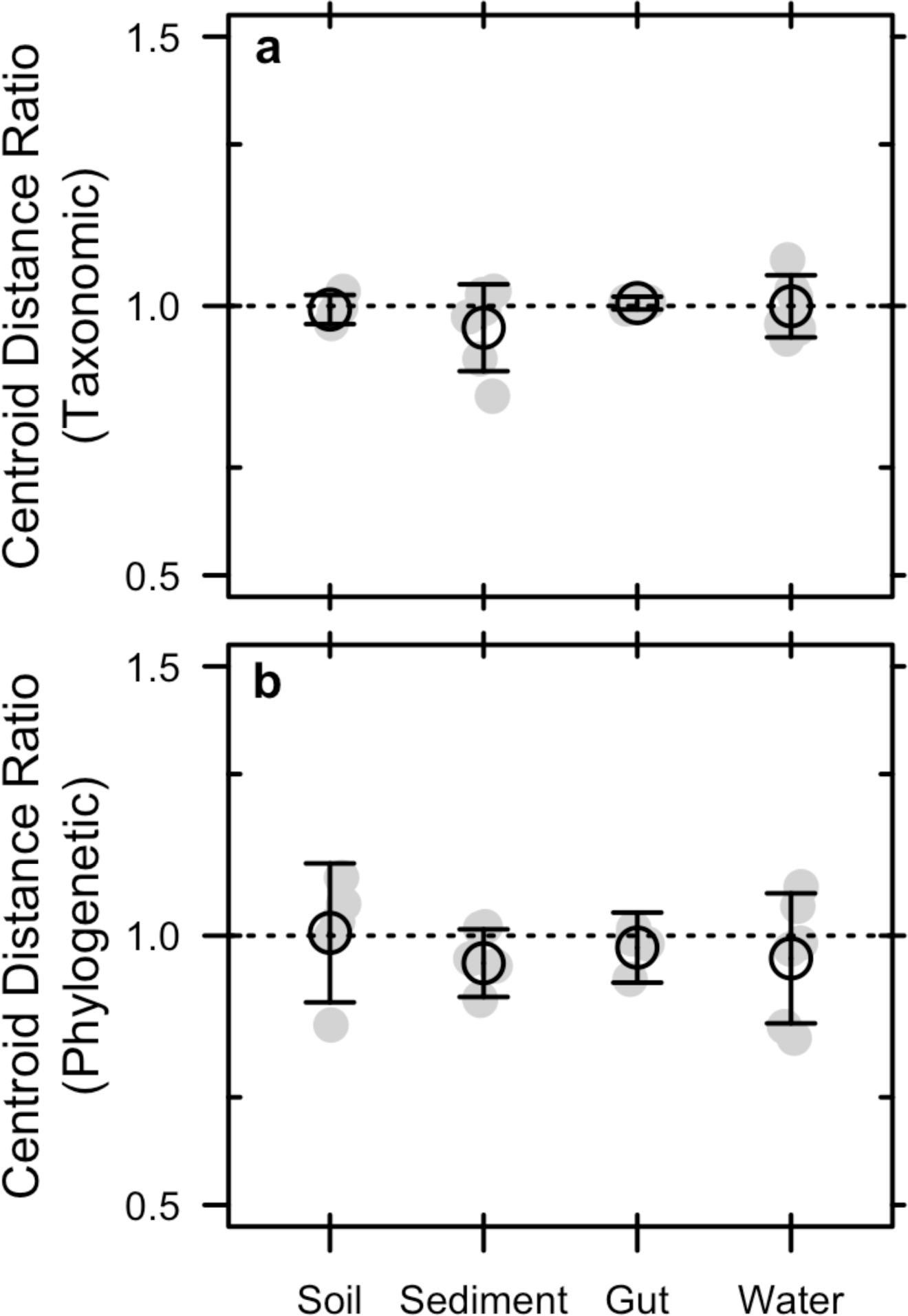
Bias of relic DNA on the among-sample bacterial diversity in different ecosystem types. **a-b** We tested for the effects of bias caused by relic DNA by calculating a beta-diversity ratio based on centroid distances. Centroid distances were estimated after performing Principle Coordinates Analyses (PCoA) using taxonomic (**a**) and phylogenetic (**b**) distance metrics (Bray-Curtis and UniFrac, respectively). The centroid distance ratio was calculated on each sample within an ecosystem type and reflects the composition of the total DNA pool (intact + relic) relative to the intact DNA pool. Relic DNA had no effect on beta-diversity for any of the ecosystem types sampled. Grey symbols are the observed data and black symbols represent means ± 95% confidence intervals.

## DISCUSSION

Rarely, if ever, are biological communities completely censused. As a result, estimates of diversity are often based on incomplete sampling, which introduces uncertainty and potential bias^25^. For example, microbiologists using cultivation-independent approaches commonly estimate the diversity of a community using hundreds of thousands of sequences from samples that contain in excess of a billion individuals. Diversity estimation of microbial communities is further complicated by the accumulation and persistence of relic DNA, a pool of sequences that may not accurately reflect the composition of viable microorganisms. We developed a set of models that capture the fundamental processes regulating relic DNA dynamics along with the effects of sampling from a joint species abundance distribution (SAD) that contain sequences from both the relic and intact DNA pools. Our models reveal that two criteria are required in order for relic DNA bias to emerge. First, the relic DNA pool must be large enough so that it diminishes the probability of sampling sequences from the intact DNA pool. Second, the relic SAD must be distinct from the intact SAD. If these conditions are not satisfied, relic DNA has minimal effect on estimates of biodiversity. We then explored the model’s expectations by analyzing the intact and total (relic + intact) DNA in ecosystems where relic DNA dynamics are known to vary. Despite making up a large portion of the total DNA, we found that relic DNA had minimal to no effect on estimates taxonomic and phylogenetic diversity, consistent with model scenarios where relic DNA degradation rates are equivalent among species.

Despite sampling a range of habitats with attributes that affect relic DNA turnover, our data suggest that estimates of taxonomic and phylogenetic diversity were unbiased by relic DNA (Figs. 2 and 3) even when it accounted for > 80 % of the total DNA (Figs. S5 and S8). Similar findings have been reported elsewhere. From a survey of North American soils, no bias was detected for taxonomic richness in 30 % of bacterial samples (7 / 31) and 55 % of fungal samples (17 / 31), even when relic DNA made up a substantial portion of the total DNA pool ^21^.Likewise, more than 90 % of the bacterial DNA recovered from porcine lung tissues was sensitive to DNase, yet diversity in the enzymatically treated and untreated samples was the same indicating that large relic DNA pools do not necessarily obscure estimates of microbial diversity ^20^. Such findings are supported by our modeling results: relic DNA has no effect on estimates of microbial diversity when the SADs of the relic and intact DNA pools are equivalent (Fig. 1b). When this assumption is met, the abundance distribution of relic and intact DNA pools should be identical. Therefore, sampling from the relic DNA pool should not bias estimates of diversity in the intact DNA pool, even if relic DNA makes up a large portion of the total DNA pool (Fig. 1b).

**Figure 3.**
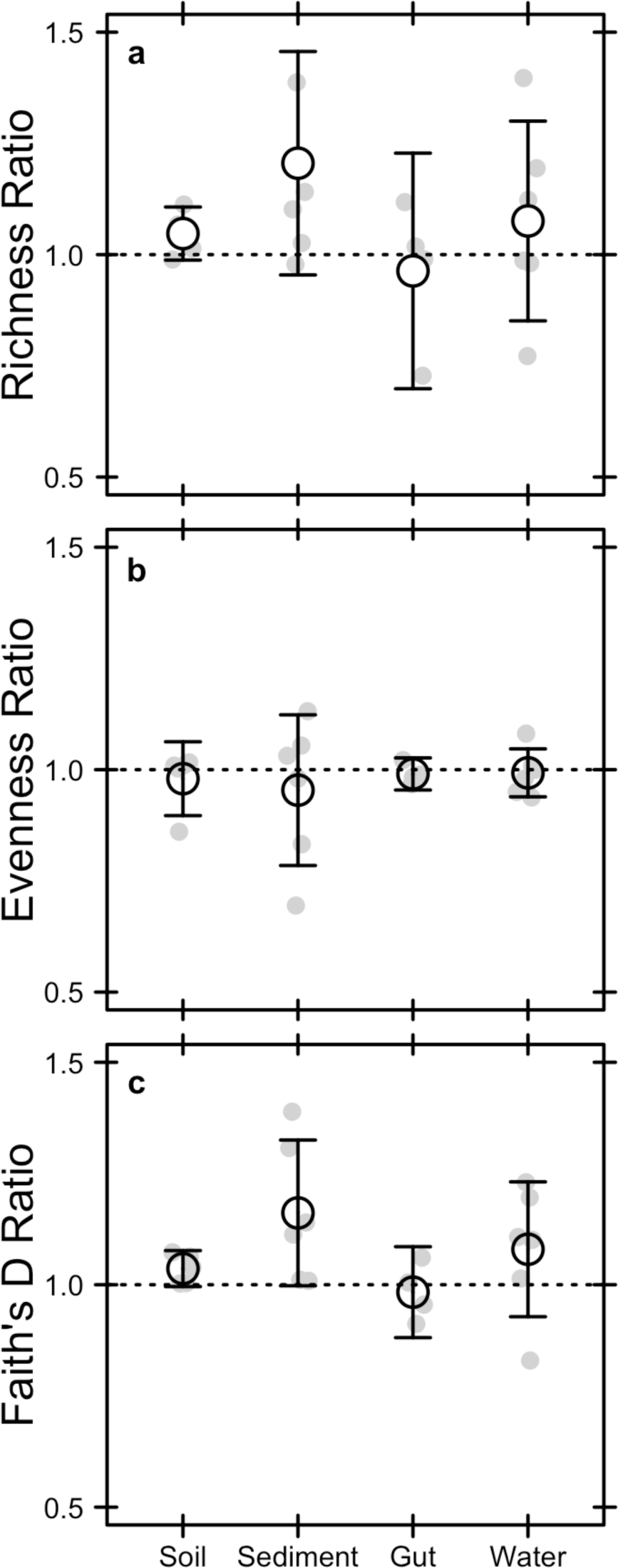
Bias of relic DNA on within-sample bacterial diversity in different ecosystem types. **a-c** We tested for the effects of bias caused by relic DNA by calculating diversity ratios for (**a**) richness, (**b**) evenness, and (**c**) phylogenetic diversity. The ratios reflect the diversity of the total DNA pool (intact + relic) divided by the diversity of the intact DNA pool. Relic DNA did not bias any measures of diversity in any of the ecosystem types. Richness was calculated as the number of operational taxonomic units (97% sequence similarity of the 16S rRNA gene), evenness was calculated using Simpson’s evenness index, and phylogenetic diversity was calculated using Faith’s *D* index. Grey symbols are the observed data and black symbols represent means ± 95% confidence intervals.

Our simulations identify other ecological scenarios where relic DNA bias may arise. It has been hypothesized that relic DNA can be protected from degradation inside of aggregates, biofilms, or other complexes that reduce contact between nucleic acids and extracellular enzymes. Importantly, protection can only increase bias if it alters relic DNA degradation in a species-specific manner. While most studies have emphasized the potential for protection to overestimate diversity, our model predicts the opposite (Fig. 1b). Protection created a less even relic SAD that resulted in the increased dominance of relic sequences associated with a few species. Assuming a large relic DNA pool, preferential sampling of these protected sequences will lead to the underestimation or richness, which has been observed in some instances ^20^. In contrast, we identified at least one ecological scenario where relic DNA creates overestimation bias. Microscale heterogeneity in structured habitats can lead to non-uniform distributions of microorganisms and their metabolic activities. Our simulations indicate that the resulting “hot spots” can degrade relic DNA sequences in a density-dependent matter resulting in a more even relic SAD. Although not significant, sediment samples trended towards positive bias for richness and PD (Fig. 3), which could potentially reflect “hot spots” of relic DNA degradation. Inflation bias can also arise when species in the regional pool disperse into habitats for which they are poorly adapted. When these immigrants die, they can enrich the relic DNA pool with sequences that are dissimilar to sequences found in the local community. A similar effect could arise when dead bacteria are transported across ecosystem boundaries, a phenomenon that occurs, for example, when marine snow is exported from surface waters to marine sediments ^26^. More generally, bias may arise under non-equilibrium conditions where community turnover of the intact DNA pool is shorter than turnover of the relic DNA pool. Any abiotic or biotic perturbation that removes a substantial amount of living biomass could result in transient divergence in the composition of sequences in the intact and relic DNA pool. For example, virulent phage can dramatically reduce the abundance of bacterial prey and in the process release a large pulse of DNA into the environment, which could bias estimates of microbial diversity ^27^. Last, although not fully explored here, shifts in rank abundance of taxa that are independent of parameters describing the SAD could also result in biased estimates of microbial diversity. In sum, there are ways to deviate from neutral expectations about the degradation of relic DNA, which should lead to biased estimates of microbial diversity, but sampling from a range of ecosystems suggests that such conditions are not prevalent in nature.

Our understanding of the microbial biosphere has been transformed by the development and application of molecular-based cultivation-independent techniques. The ability to rapidly obtain millions of gene sequences and transcripts from a range of environments has yielded valuable insight into the processes that regulate community assembly and function ^28,29^, and has also paved the way for the discovery of new metabolisms ^30^, tests for unifying patterns of biodiversity ^31^, and an updated tree of life ^32^. There are limitations, however, associated with culture-independent techniques, which include inefficient nucleic-acid extraction methods ^32^ and “universal” primers that over-represent some taxonomic groups while overlooking others ^33^. Sequencing of relic DNA is another important concern, which can potentially lead to the overestimation or underestimation of microbial diversity. However, this bias requires the decoupling of processes that regulate the compositional turnover of the relic and intact DNA pools. While some recent evidence suggests this can arise ^20^, our findings suggest that at least in some ecosystems, relic DNA appears to contribute minimally to the characterization of microbial community structure.

## METHODS

### Sampling Model

We used a set of sampling-based simulations to explore the effects of mixing intact and relic DNA on estimates of diversity. For each simulation, we defined a regional species pool consisting of 10,000 taxa with a lognormal abundance distribution ^34^. The intact community consisted of 1,000,000 individuals sampled from this regional pool. We then combined this intact community with a relic community at proportions ranging from 0.01 to 0.96. We altered the evenness of the regional pool from which the relic community was sampled by changing the scale parameter of the lognormal distribution. To decrease the evenness of the relic DNA pool, we increased the scale parameter from 0.98 to 1.8. To increase the evenness of the relic DNA pool, we decreased the parameter from 0.98 to 0.25. After mixing the intact and relic communities at the defined proportions, we rarified the total community to 10,000 observations and calculated richness and Bray-Curtis distances to estimate compositional differences between the intact and total DNA pools. To estimate contribution of relic DNA to diversity, we calculate ratios (total DNA / intact DNA) for richness and Bray-Curtis distances. All simulations and estimations were performed in the R statistic environment (v 3.3.2)^35^ using the ‘vegan’ package as well as custom functions.

### Process-Based Model

We developed a set of stochastic simulations to explore how the processes regulating relic DNA dynamics can give rise to bias in the estimation of microbial diversity. For each simulation, we defined a regional species pool consisting of 4,000 taxa with a lognormal abundance distribution ^34^. We initiated an intact community by randomly sampling 20,000 individuals from the regional species pool. Subsequent dynamics were controlled by the processes depicted in our conceptual model (Fig. 1a). Specifically, we simulated immigration by sampling *j* individuals from the regional species pool and adding them to the intact community. We simulated birth by randomly selecting *I × r* individuals from the intact community and adding them to the intact community again, where *I* is the size of the intact community and *r* is the per capita birth rate. We simulated death by selecting (*I* × *m*) + *j* individuals from the intact community and moving them to the relic community (*R*), where *m* is the per capita mortality rate. We simulated degradation by selecting (*R* × *d*) + *j* individuals from the relic community and removing them, where *d* is the per capita decay rate of relic DNA.

We simulated three scenarios with the process-based model. First, we considered a neutral scenario. At each time step, we simulated degradation by randomly selecting relic DNA sequences and removing them. Second, we considered the scenario where some relic DNA sequences have lower rates of degradation owing to protection. Each species was randomly assigned a degradation susceptibility probability from a beta distribution. The beta distribution is a continuous probability distribution bound by 0 and 1 with two shape parameters. We chose shape parameters (α = 0.7, β = 0.7) to provide a wide range in susceptibility probability but also have increased probabilities of low (near 0) and high (near 1) susceptibility, which created a wide U-shaped distribution. Susceptibility probabilities were randomly assigned to species for each model iteration and were used to weight the probability of selecting relic DNA sequences for degradation at each time step. Last, we considered the scenario where there are “hot spots” of relic DNA degradation, reflecting the clumped distribution of microbial taxa and their associated metabolic activities. At each time step, we calculated density-dependent selection probabilities by weighting the probability of selecting an individual from the relic community by its species abundance. Therefore, relic DNA sequencing belonging to more abundant species would be more likely to be selected for degradation. For each of the three scenarios, we ran each simulation for 10,000 time-steps. We used a constant immigration rate (*j* = 2,000) to maintain intact community diversity, and used equal and constant birth and mortality rates (*b* and *m* = 0.1) to maintain intact community size. Likewise, we accounted for immigration during death and degradation to prevent uncontrolled growth. For each set of simulations, we used a range of decay rates (*d*), which yielded relic DNA proportions between 0.05 and 0.95 according to the following equation: *d* = *m/p* − *m*, where *p* is the target proportion and *m* is the fixed mortality rate (see Supplement for derivation). At the end of each simulation, we determined the effect of relic DNA on diversity estimates by comparing the total community (intact + relic) with the intact community. All simulations were performed in the R statistic environment.

### Sample collection and DNA pools

We collected samples from a diverse set of environmental and host-associated ecosystems. First, we sampled sediments and surface water from lakes near the Michigan State University, W.K. Kellogg Biological Station (KBS) in Hickory Corners, Michigan, USA. Soils were sampled from the main sites and surrounding areas at the KBS Long-Term Ecological Research site ^36^. We also collected fresh feces as representative gut samples from cows, dogs, horses, rabbits, and humans. In each of these ecosystem types, we obtained samples from 6 - 8 independent sites. In the laboratory, we applied a DNase treatment to each sample to remove relic DNA. The procedure is based on methods that have previously been used to quantify relic DNA in marine sediments ^37,38^, host tissue ^37,38^, and drinking-water biofilms ^37,38^. The DNase-based method has proven to be effective not only for removing extracellular DNA, but also DNA that is contained inside of dead cells ^37,38^ while not compromising the integrity of living cells ^37,38^. We determined that the DNase method is efficient (98%) at removing relic DNA that was experimentally added to an environmental sample (see Supplementary Methods; Fig. S4). Thus, we refer to the DNA remaining in a subsample following enzymatic treatment as “intact DNA” and assume it is derived from viable cells. “Total DNA” refers to the DNA in subsample that was not treated with DNase (negative control), which reflects the sum of intact DNA and relic DNA.

For aquatic samples, we filtered 250 mL lake water through a 47 mm 0.2 μm Supor PES membrane filter using 10 mm Hg vacuum. We cut the filter in half and randomly assigned one half of the filter to a DNase treatment and used the other half as the control. We then rolled up each piece of filter using sterile forceps and inserted it into a 2 mL centrifuge tube containing 1.5 mL of pH 7.3 phosphate buffered saline (PBS). After vortexing at room temperature for 5 min, we removed the filter from the tube and centrifuged for 5 min at 10,000 x g. We then discarded the supernatant and resuspended the pellet in 375 μL of PBS, which was transferred to a 2 mL centrifuge tube. For non-aqueous samples (soil, sediments, and feces), we directly added 0.25 g of material to a 2 mL centrifuge tube. At this stage in the procedure, aquatic and non-aquatic samples were identically processed. For each sample in its 2 mL centrifuge tube, we added DNase digestion buffer, which consisted of 382.5 μL of nanopure water, 5 μL of 1 M MgCl_2_, 2.5 μL of bovine serum albumin (10 mg/mL), and 50 μL of 0.5 M Tris HCl (pH 7.5). For subsamples treated with DNAse, we added 40 μL of a 10 U/μL stock of DNase I (Roche # 04536282001) and 20 μL of nanopure water resulting in a 500 μL final working volume with an 80U/μL DNase concentration. (See Supplementary Material and Fig. S3 for description of experiment to determine minimum DNase concentration needed for relic DNA removal.) For untreated subsamples, we substituted 40 μL of nanopure water for DNase solution. For each sample, we measured pH using a micro pH probe (Orion 9110DJWP, Thermo Scientific) and adjusted to 7.3 - 7.7, which is in the optimum range for DNase. We then incubated the samples horizontally on a shaker table at 37 °C for 60 min. Following this, we transferred each sample to a 15 mL Falcon tube containing 1 mL of 1X hexadecyltrimethylammonium bromide (CTAB) buffer, which consisted of one part solution 1 (1 g CTAB + 0.58 g NaCl in 10 mL of nanopoure water) and one part solution 2 (0.58 g NaCl + 0.82 K_2_HPO_4_ + 0.04 KH_2_PO_4_ in 10 mL of nanopoure water). We then added 25 μL of 0.5 M EDTA to each tube and vortexed. We stopped the DNase reaction by incubating the tubes at 75 °C for 10 min. We began DNA extraction by adding 1 mL of phenol:chloroform:isoamylalcohol (25:24:1) to the 2 mL tube, which was then vortexed for 10 min. This was followed by centrifuging at 7,100 x *g* for 10 min. We transferred the top aqueous layer to a 15 mL Falcon tube and combined it with an equal volume of chloroform:isoamylalcohol (24:1) and then vortexed. After centrifuging the Falcon tubes at 7,100 x *g* for 5 min, we transferred 400 μL of the top aqueous layer to a new 2 mL centrifuge tube. For clean up, we used the MO BIO PowerLyzer PowerSoil DNA Isolation Kit starting from Step 9. At Step 21 of the DNA Isolation Kit, we eluted in 50 μL of Solution C6 and centrifuged for 1 min at 10,000 x g. We stored extracted DNA at −20 °C for short-term storage or at −80 °C for long-term storage.

### Contribution of relic DNA to bacterial abundance

We used 16S rRNA gene copy numbers generated from quantitative PCR (qPCR) assays to estimate the proportion of relic DNA in a sample as 1 - (intact DNA / total DNA). Briefly, the previously described qPCR assays^39^ consisted of 30 μL reactions containing 1μL of DNA template, 0.5 μL of each primer (10 μmol/L), 14.5 μL of nuclease-free H_2_O, and 13.5 μL of 5 Prime 2.5x RealMasterMix SYBR ROX. We amplified a 200 base-pair fragment of the 16S rRNA gene with Eub 338 (forward) and Eub518 (reverse) primers ^40^. PCR assays were performed with an Eppendorf Mastercycler Realplex^2^ system using previously reported thermal cycle conditions ^40^. We generated qPCR standards from bacterial genomic DNA (*Micrococcus* sp.) using the TOPO TA Cloning Kit (Invitrogen). We extracted plasmids from transformed cells ^41^, and used the M13 forward and reverse primers to generate PCR products. The PCR products were quantified and used to generate a standard curve capturing a range of 10^2^ - 10^7^ gene copies/μL. We used a noiseband threshold for quantification. The coefficients of determination (r^2^) for our assays ranged from 0.96 and 0.99, while amplification efficiencies fell between 0.93 and 0.99. Based on melting curve analyses, we found no evidence for primer dimers. All unknown samples, no template controls, and standards were run in triplicate on every plate. The mean coefficient of variation (standard deviation / mean) for our 16S rRNA qPCR assay was 0.16.

### Contribution of relic DNA to bacterial diversity

#### Community sequencing

We estimated the contribution of relic DNA to bacterial diversity using high-throughput sequencing of the 16S rRNA gene. Specifically, we amplified the V4 region of the 16S rRNA gene from the intact and total DNA pools of each sample using barcoded primers (515F and 806R) designed to work with the Illumina MiSeq platform ^42^. We cleaned the sequence libraries using the AMPure XP purification kit, quantified the resulting products using the QuantIt PicoGreen kit (Invitrogen), and pooled libraries at equal molar ratios (final concentration: 20 ng per library). We then sequenced the pooled libraries with the Illumina MiSeq platform using paired end reads (Illumina Reagent Kit v2, 500 reaction kit) at the Indiana University Center for Genomics and Bioinformatics Sequencing Facility. Paired-end raw 16S rRNA sequence reads were assembled into contigs using the Needleman algorithm ^43^. We obtained a total of 12,916,632 16S rRNA sequences. After quality trimming with a moving average quality score (window 50 bp, minimum quality score 35), we aligned the sequences to the Silva Database (version 123) using the Needleman algorithm. Chimeric sequences were removed using the UCHIME algorithm ^44^. After this filtering, there was an average (± SEM) of 222,701 ± 9,560 sequences per site. We created operational taxonomic units (OTUs) by first splitting the sequences based on taxonomic class (using the RDP taxonomy) and then binning sequences into OTUs based on 97% sequence similarity. Our depth of sequencing led to a high degree of coverage across samples (minimum Good’s Coverage = 0.98). For phylogenetic analysis, we picked representative sequences for each OTU by using the most abundant unique sequence. We used FastTree ^45^ to generate a phylogenetic tree from the representative sequences using the generalized time-reversible model of nucleotide evolution. We calculated phylogenetic distances using weighted UniFrac distances ^46^. All initial sequence processing was completed using the software package mothur (version 1.38.1) ^47^.

#### Alpha Diversity

We estimated the effects of relic DNA on richness, evenness, and phylogenetic diversity for the intact and total DNA pools with a sample. To estimate the number of OTUs (richness), we used a resampling approach that subsampled each sample to an equal number of sequences per sample and summed the number of OTUs that were represented ^48^. Briefly, we subsampled to 30,000 observations, resampled 999 additional times, and then calculated the average richness estimates (± SEM) for each sample. To estimate the equitability in abundance among taxa in a sample (evenness), we used the same resampling approach and calculated average evenness estimates (± SEM) using Simpson’s Evenness index ^49^. To test whether relic DNA affected the phylogenetic diversity within a sample, we subsampled communities to 30,000 observations and then calculated Faith’s *D* statistic, which sums the branch lengths for each species found in a sample from the root to the tip of the phylogenetic tree ^50^. All estimations were performed in the R statistic environment (v 3.3.2)^35^ using the vegan, ape, ade4, picante, and plyr packages, along with custom functions.

#### Beta Diversity

We estimated the effects of relic DNA on between-sample (beta) diversity by comparing the taxonomic and phylogenetic diversity of bacterial communities in the intact and total DNA pools. First, we conducted a Principal Coordinates Analysis (PCoA) on log_10_-transformed relative abundance data to visualize the effects of relic DNA removal (via DNase treatment) on bacterial community composition within and among ecosystem types. The PCoA was performed with Bray-Curtis and UniFrac distances to assess taxonomic and phylogenetic effects, respectively. In addition, we used PERMANOVA to test for differences in taxonomic and phylogenetic composition based on ecosystem type for the total DNA pool. Second, we conducted a Mantel test to assess the correlation between the community resemblance matrices (either Bray-Curtis or UniFrac) represented by the intact and total DNA pools. Last, we tested whether relic DNA altered beta-diversity within an ecosystem type by comparing centroid distances. To calculate this metric of sample dispersion, we determined the centroid from a PCoA with either Bray-Curtis or UniFrac distances for the total DNA pool for all sites within a given ecosystem type. We then measured the Euclidean distances between the centroid and all samples (total and intact) to determine the centroid distances (see Fig. S7).

## ACKNOWLEDGMENTS

We acknowledge constructive feedback from KD Webster, RZ Moger-Reischer, EK Hall, NI Wisnoski, and WR Shoemaker. We thank S. Kuenzel and B.K. Lehmkuhl for technical support. This work was supported by the USDA National Institute of Food and Agriculture Grant 2011-67019-3022 (JTL), the National Science Foundation Dimensions of Biodiversity Grant 1442246 (JTL), and US Army Research Office Grant W911NF-14-1-0411 (JTL). All code and data used in this study can be found in a public GitHub repository (https://www.github.com/LennonLab/relicDNA) and the NCBI SRA.

